# Puerarin blocks aging phenotype in cultured human dermal fibroblasts

**DOI:** 10.1101/2020.09.29.319616

**Authors:** Yuki Kamiya, Mao Odama, Aki Mizuguti, Shigeru Murakami, Takashi Ito

## Abstract

Dermal fibroblast aging contributes to aging-associated functional defects in the skin since dermal fibroblasts are important to maintain skin homeostasis by interacting with epidermis and extracellular matrix. Here we identified that puerarin, an isoflavone contained in *Pueraria lobata* (Kudzu), can prevent the aging-phenotype of human dermal fibroblasts. Puerarin treatment increased in proliferating cells and decreased in senescence-associated beta-galactosidase positive cells in the high-passage culture of dermal fibroblasts. Moreover, puerarin reduced smooth muscle actin-positive myofibroblasts and the expression of a reticular fibroblast marker, calponin 1 (CNN1), which were induced in high-passage fibroblasts. Fulvestrant, an estrogen receptor antagonist, blocked puerarin-mediated downregulation of SMA and CNN1. Our results suggest that puerarin may be a useful food factor that alleviates aging-related functional defects in the skin.

## Introduction

Skin aging is dependent on the diet and nutrition as well as the other intrinsic factors, such as chronological aging, and the extrinsic factors, such as sun exposure [1,2]. Since excessive reactive oxygen species (ROS) generated by the harmful stresses cause cellular senescence and aging phenotype, anti-oxidative rich foods have been well investigated as a preventive strategy against skin aging [2–4]. Additionally, recent evidence suggests that targeting aging-associated changes by natural products can prevent age-related phenotype.

Cellular senescence is a hallmark of normal aging, which is a state of irreversible growth arrest triggered by both endogenous stress and exogenous stress [5–7]. Senescent cells are characterized by the induction of senescence-associated beta-galactosidase activity, telomere shortening, the expression of cyclin-kinase inhibitors, such as p16 or p21, etc. Senescent cells secrete proinflammatory cytokines, growth factors, and matrix metalloproteases, that contributes to local and systemic dysfunction [8,9]. Accumulating evidence demonstrated that clearance of senescent cells prevents the age-associated decline of tissue function and extends lifespan [10]. Recently, small molecules, including the nutritional agents, that can selectively eliminate senescent cells, called senolytics, are expected to prevent age-associated decline of tissue function[7,11,12].

Fibroblasts play an important role in maintaining tissue structure by the synthesis of extracellular matrix proteins, such as collagen and elastin, in most tissues. The skin dermis has three layers, the papillary dermis, the reticular dermis and the hypodermis [13]. The papillary dermis is the superficial layer that contacts the epidermis and has a higher cell density, while the reticular dermis is the deeper layer and has a thicker extracellular matrix and a lower cell density [14,15]. The fibroblasts residing in papillary and reticular dermis have distinct identities. Papillary fibroblasts exhibit a spindle-shaped morphology and show higher proliferative activity than reticular fibroblasts. Meanwhile, reticular fibroblasts exhibit a polygonal-morphology and express the myofibroblast marker alpha-smooth muscle actin [16]. Aging mainly affects the cellular function of papillary fibroblasts, including a decrease in proliferative capacity and an increase in transdifferentiation to reticular fibroblasts, which may lead the aging-associated structure changes [17–20]. Recent studies showed that diet and nutrition can influence the skin phenotype by controlling the aging of dermal fibroblast [21–23], proposing that nutritional agents that modify the fibroblast function links skin aging.

Puerarin (daidzein-8-C-glucoside) is a major isoflavone found in *Pueraria lobata* (Kudzu, Kuzu, Gegen) which is an edible legumes [21–24]. Puerarin is abundant in vine and root of *Pueraria lobata*; 0.1 ∼ 1 g in 100 g of the vine, 3 ∼ 10 g in 100g of the root [25–28]. The root of *Pueraria lobata* is also used as a component of traditional Chinese medicine to treat chills, colds, fever, and diarrhea. Recent studies revealed a variety of pharmacological actions of puerarin, such as vasodilatation, cardioprotection, anticancer, menopausal symptoms, etc. [29,30]. Oral ingested puerarin is absorbed from the gut into blood; the oral bioavailability of puerarin has been reported to be approximately 4-7% in animal models [31–34]. Concerning the effects on aging, puerarin reduces the endothelial progenitor cell senescence [35]. However, the effects of puerarin on skin aging and fibroblast phenotype remain to be elucidated.

In the present study, we investigated the cytoprotective and anti-aging properties of puerarin in the normal human dermal fibroblasts in vitro.

## Methods

### Cell culture

Normal human dermal fibroblasts (PromCell, Heidelberg, Germany) were cultured in DMEM containing 10% fetal bovine serum and antibiotics (penicillin and streptomycin, Nacalaitesque Kyoto, Japan) according to the previous report [17]. Cells were subcultivated with 1:2 to 1:5 split ratio every week. High-passage fibroblasts (25-35 passages) were confirmed to have lower proliferative activity and were used as senescent cells. Low-passage fibroblasts (5-15 passages) were used as young control. Before experiments, NHDFs were plated on the 96-well plate or 6-well plate, then were treated with DMSO (1:1000) or puerarin (25-50 μM) and Fulvestrant (1μM, [36]) next day, and were cultured for more 72 h.

### Cell viability and cell injury assays

NHDFs were plated on the 96-well plate at 2,500 - 10,000 cells per well, and then were cultured with puerarin. Cell viability and cell toxicity were evaluated by Cell counting kit-8 assay and lactate dehydrogenase (LDH) assay (Dojindo, Kumamoto, Japan), respectively, according to the manufacturer’s protocols.

### Beta-Garactosidase activity

The Senescence-associated (SA) beta-galactosidase activity was detected as previously reported [37]. In brief, after fixed by 10% formalin in PBS, cells were incubated in X-gal stain solution (5mM potassium ferricyanide, 5mM potassium ferrocyanide, 2mM MgCl_2_ and 0.5 mg/mL X-gal in citric acid/potassium phosphate buffer (pH 6.0)) at 37 °C overnight. The numbers of the stained cells per microscope field of view (x100) were calculated as the senescent cells.

### BrdU assay

BrdU assay was performed as previously reported [38] with brief modification. NHDFs were plated on the 96-well plate at 5,000 cells per well. After cultured with or without puerarin, BrdU was added to the culture medium and the cells were cultured for more 24 h. After fixed by 10% formalin in PBS and then incubated with 2 mol/L HCl for 10 min at room temperature and for 20 min more at 37°C to denature DNA, BrdU incorporated in DNA was detected by the immunocytochemical method. The numbers of BrdU-positive cells per microscope field of view (x100) were calculated.

### Measurement of intracellular ROS

After cells on the 96-well plate were cultured with or without puerarin for 72 h, CellROX Green regent (final concentration: 5μM, ThermoScientific, USA) was added into the culture medium, and incubated for more 30 min. Then cells were washed and fixed with 10% formalin in PBS, and cells were observed by fluorescent microscopy. The fluorescent intensity was determined by using ImageJ software.

### mRNA measurement

Total RNA was isolated from NHDFs by using FastGene™ RNA Basic Kit (Fast Gene, Japan) and cDNA was generated by using Rever Tra Ace (Toyobo, Osaka, Japan). Quantitative RT-PCR analysis was performed by using Applied Biosystems Step One (Applied Biosystems) with THUNDERBIRD SYBR qPCR Mix (Toyobo) according to the previous report [39]. The primers used are shown in Table 1. To consider the specificity of the primer set for SMA, a set of primers which has been designed to selectively detect the human SMA was used [40]. The primers for the other genes were originally designed by using Primer-BLAST tool (http://www.ncbi.nlm.nih.gov/tools/primer-blast/).

**Table 1.**
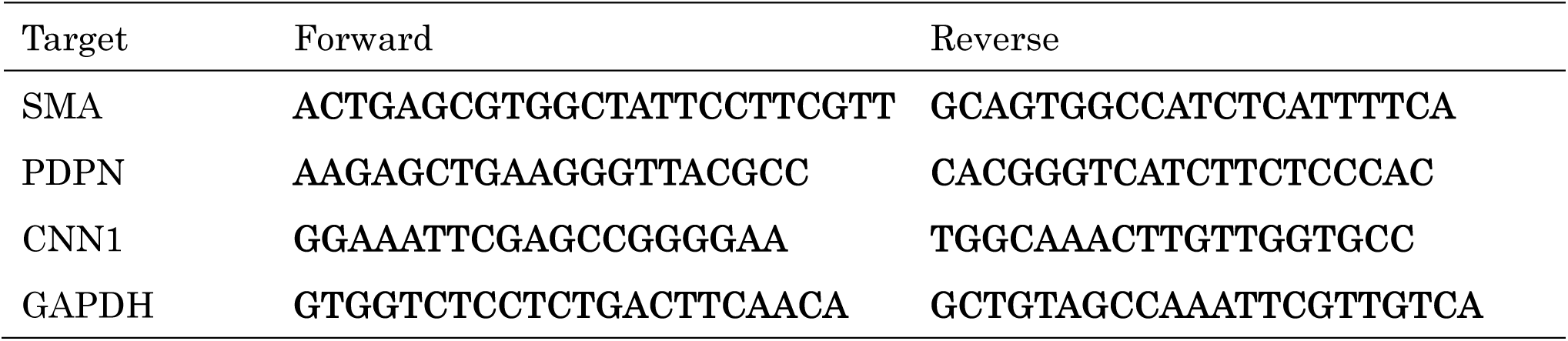
Primers used for real-time PCR

### Immunofluorescence microscopic examination

After NHDFs were fixed by 10% formalin in PBS and then were permeabilized by 0.1 % Triton-X 100 in PBS for 10 min, and blocked in BlockingOne regent (Nacalai tesque, Kyoto, Japan). Immunostaining was performed by using specific primary antibodies and the fluorescent secondary antibodies diluted in Can Get Signal Immunostain (Toyobo) according to the previous report [41]. Nuclear were stained with DAPI. Cells were examined by Fluorescent microscopy (Leica). Antibodies used in this study are as follows; Mouse anti-α-SMA actin IgG (Abcam), Rat anti-BrdU IgG (Abcam), Alexa 555-conjugated goat anti-mouse IgG antibody (Thermoscientific), CF488-conjugated anti-rat IgG antibody (Biotium).

### Statistical analysis

Student’s t-test or t-test with Bonferroni correction (for multiple comparisons) were used to determine statistical significance between groups. Differences were considered statistically significant when the calculated P-value was less than 0.05.

## Results

### The protective effects of puerarin in senescent NHDFs

To explore the effect of puerarin against skin aging, we tested the effect of puerarin on the cell viability in replication-induced senescence (Fig. 1). After passing (25∼35 passages), the proliferation of NHDFs became slow. Treatment of puerarin (25-50 μM) for 72h increased the cell viability of high passaged senescent NHDFs, although it did not influence in low passaged young NHDFs (Fig. 1A,B). Cell injury during normal culture, as measured by LDH release, was attenuated by puerarin treatment in senescent NHDFs (Fig. 1C). We investigated the effect of puerarin on the proliferation rate in high-passaged NHDFs, as assessed by BrdU assay (Fig. 1D,E). BrdU positive cells in microscopic fields were increased in puerarin-treated NHDFs, indicating that puerarin enhances cell proliferation.

**Fig. 1.**
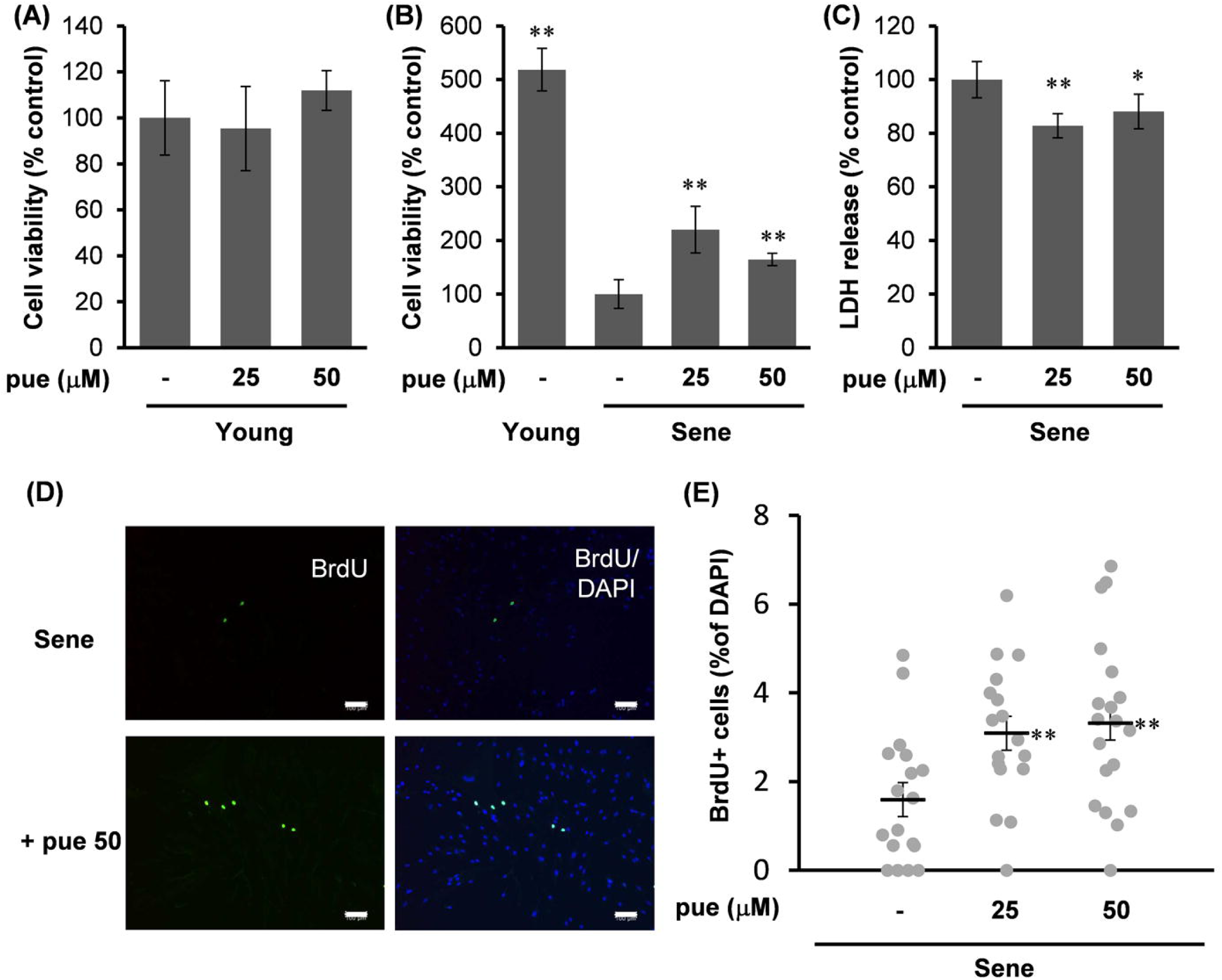
The effect of puerarin on the viability, injury and proliferative activity in high-passage NHDFs. A, B: The effect of puerarin (pue) on cell viability of young (A) and senescent NHDFs (B) was analyzed by CCK-8 assay. n=5. C: The effect of cellular LDH release in high passage NHDFs. Data was obtained from LDH assay. n=6. D, E: The effect of puerarin on the cell proliferative activity in senescent NHDFs. Cells were stained for BrdU and nuclei by specific antibody and DAPI, respectively, and photographed at 100 × magnification (D). The percentage of BrdU positive cells per total cells was calculated from 15 microscopic fields for each group. (E). Scale bars = 100μm. *; p<0.05, **; p<0.01 vs control.

Moreover, we evaluated the effect of puerarin on the ROS generation in senescent NHDFs, as assessed by the fluorescent ROS probe (Fig. 2). Since isoflavones possess an anti-oxidative role [36], we expected that puerarin lowers ROS production. However, puerarin treatment rather increased ROS production in senescent NHDFs.

**Fig. 2.**
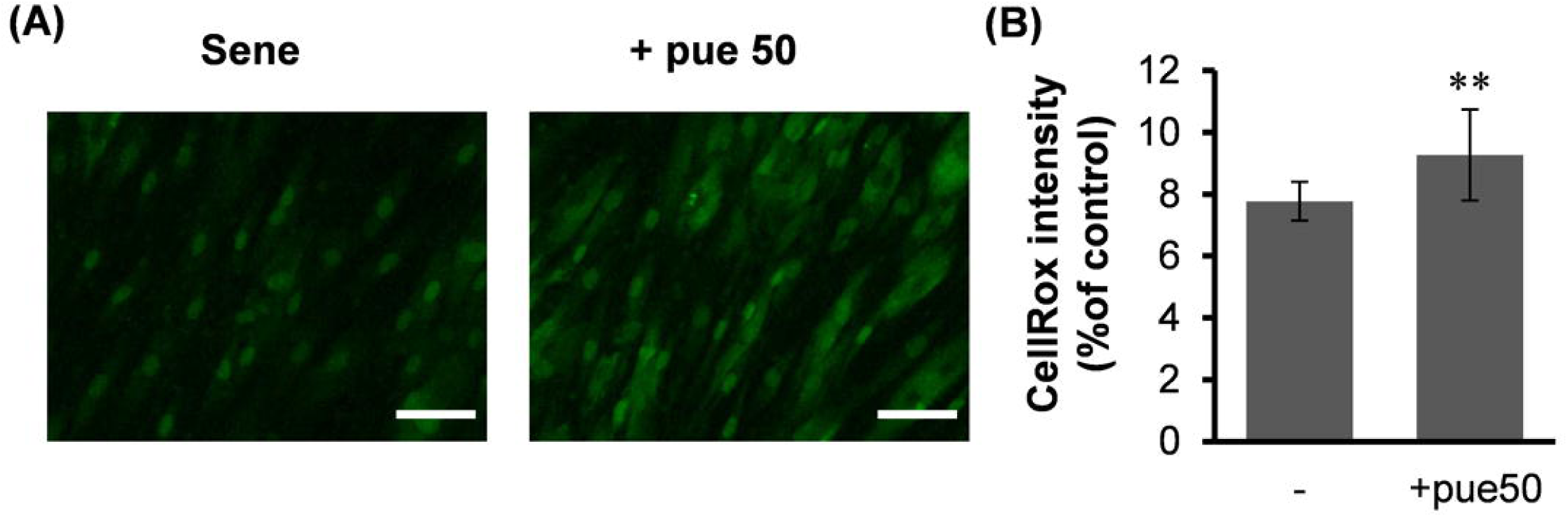
The effect of puerarin on ROS generation in high-passage NHDFs. ROS generation in senescent NFDFs (sene) with or without 50 μM puerarin (+pue 50) was measured by fluorescent ROS probe CellROX Green (A). CellROX intensity was calculated from 10 images for each group (B). Scale bars = 100μm. **; p<0.01 vs control.

### The anti-aging effects of puerarin in senescent NHDFs

Next, we evaluated the effect of puerarin on the senescent phenotype of NDHF. SA-beta-galactosidase positive NDHFs were decreased by puerarin treatment (Fig. 3A,B).

**Fig. 3.**
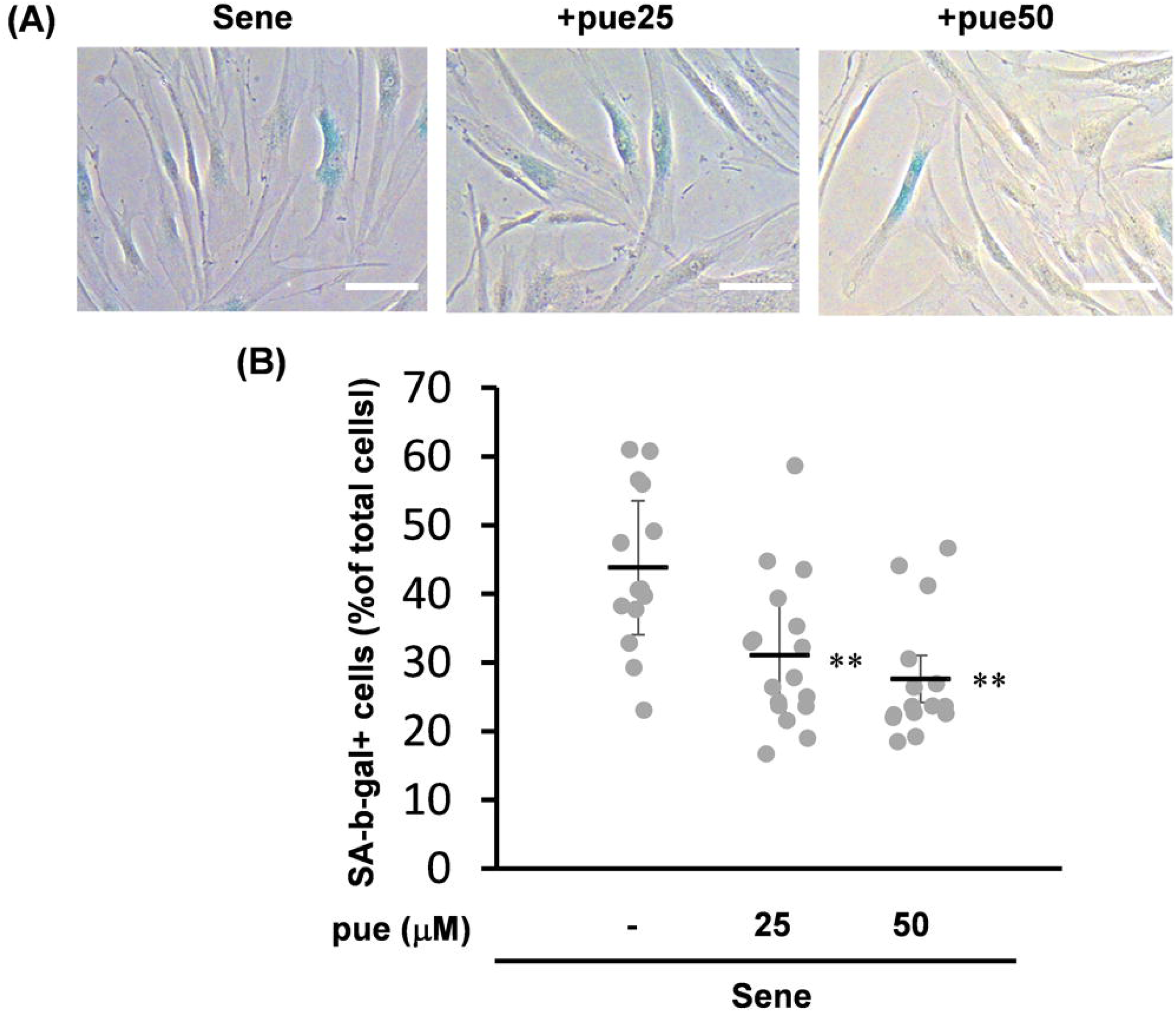
The effect of puerarin on cellular senescence in high-passage NHDFs. A, B: The effect of puerarin on SA-β-galactosidase activity. Young and senescent (sene) NHDFs treated with 20 or 50 μM puerarin (pue) were stained by SA-β-galactosidase assay (A). The percentage of SA-β-galactosidase positive cells per total cells was calculated from 15 microscopic fields for each group. Scale bars = 100 μm. *; p<0.05, **; p<0.01 vs control.

Another characteristic of aged dermal fibroblasts is a decrease in papillary fibroblasts and an increase in the reticular fibroblasts in the skin [18,22]. In the case of the NHDF culture model, high passage of NHDFs causes an increase in the cells of reticular phenotype and smooth muscle actin (SMA)-positive myofibroblasts [19,42]. Therefore, we first evaluated the effect of puerarin on the frequency of α-SMA-positive cells. Puerarin treatment reduced the numbers of α-SMA-positive cells in high passaged senescent NHDFs (Fig. 4A). A reduction in SMA by puerarin treatment was also validated by mRNA expression (Fig. 4B).

**Fig. 4.**
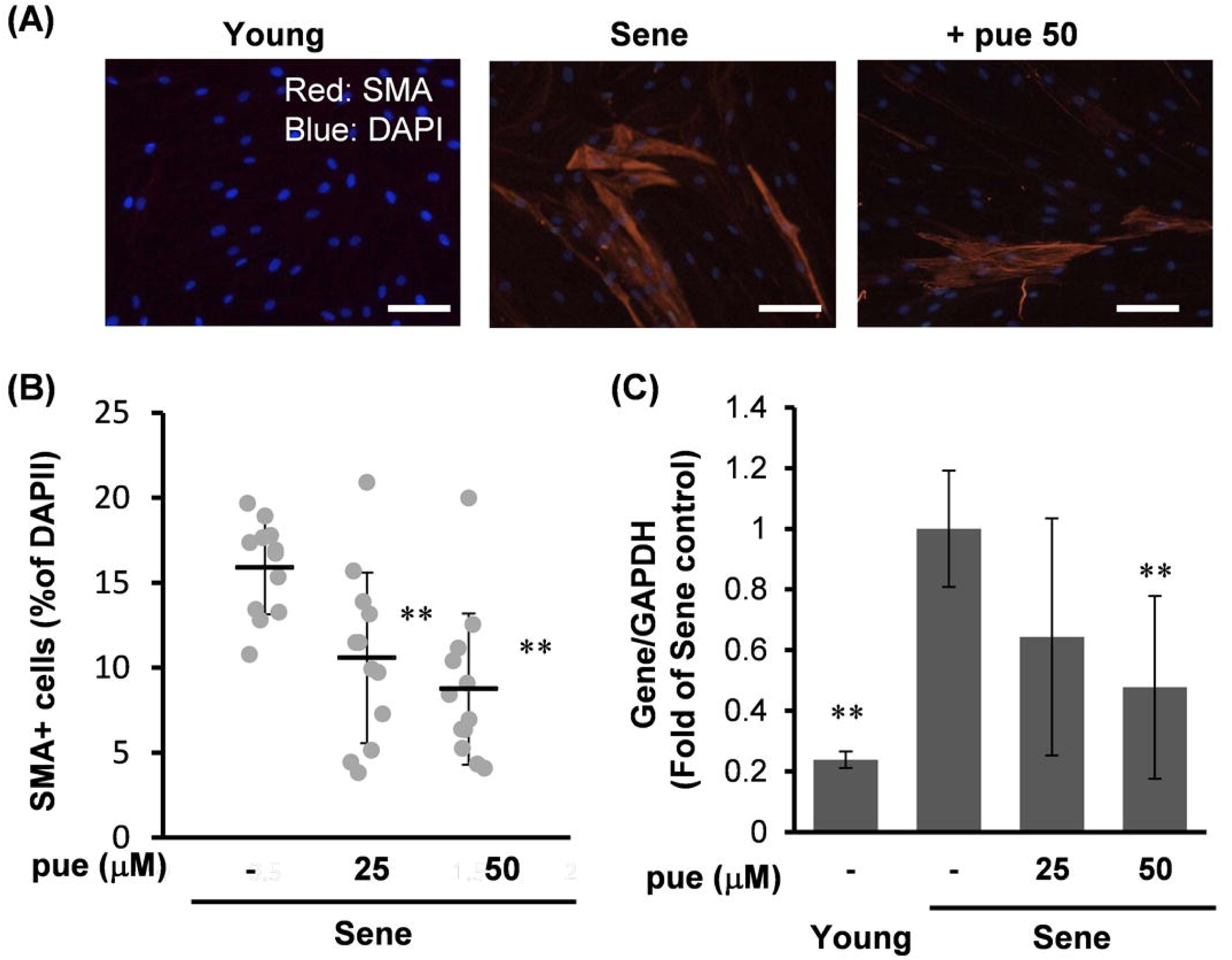
The effect of puerarin on the expression of SMA in high passage NHDFs. A-B: Young or senescent (sene) NHDFs with or without 50 μM puerarin (+pue 50) were stained for SMA by the immunocytochemical method. The percentage of SMA positive cells per total cells was calculated from 15 microscope fields for each group. C The expression of SMA mRNA in young and senescent NHDFs treated with or without 25-50 μM puerarin (pue) for 72 h. Scale bars = 100μm. n=4 *; p<0.05, **; p<0.01 vs control.

Next, we evaluated the effect of puerarin on the gene markers of papillary and reticular phenotype, podoplanin (PDPN), and calponin-1 (CNN1), respectively (Fig. 5). Puerarin treatment upregulated PDPN expression in senescent NHDFs. In contrast, puerarin treatment reduced CNN1 expression in senescent NHDFs, while the CNN1 expression was increased by high passage compared to low passage NHDFs, indicating that puerarin prevents the transdifferentiation to reticular fibroblasts.

**Fig. 5.**
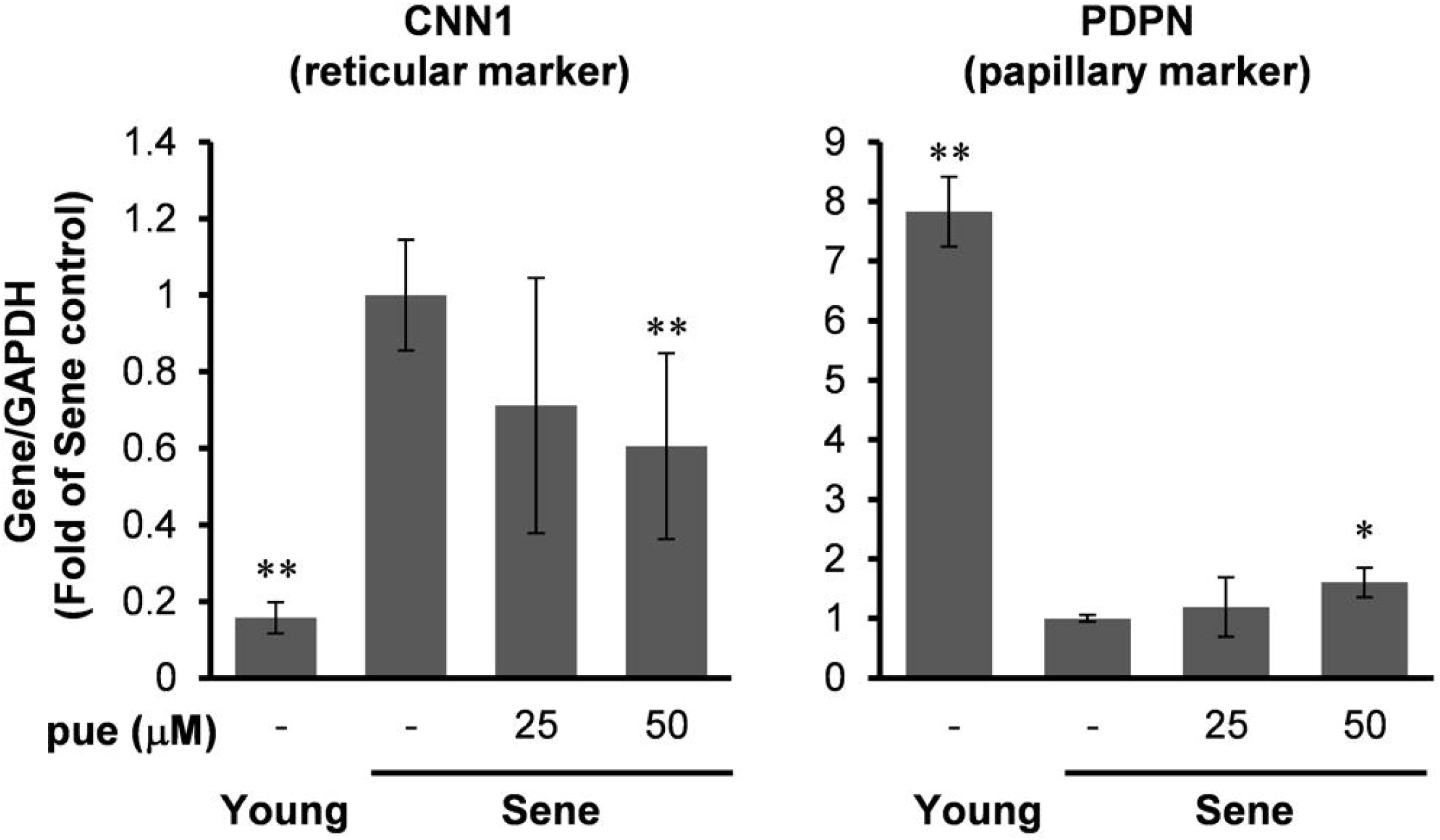
The effect of puerarin on gene expressions of papillary and reticular markers. The expressions of PDPN (papillary marker), CNN1 (reticular marker) in young and senescent NHDFs treated with or without 25-50 μM puerarin (pue) for 72 h. n=3-5 *; p<0.05, **; p<0.01 vs control.

### The role of Estrogen receptor signaling pathway on the anti-aging effects of puerarin

Since isoflavones possess an estrogen receptor (ER) agonist property, we tested whether the ER pathway is involved in the effect of puerarin on the gene expression in NHDFs. An ER antagonist, Fulvestrant, blocked puerarin-dependent downregulation of SMA and CNN1 (Fig. 6). These data indicate that the attenuation of transdifferentiation to reticular fibroblasts by puerarin may be associated with the ER pathway.

**Fig. 6.**
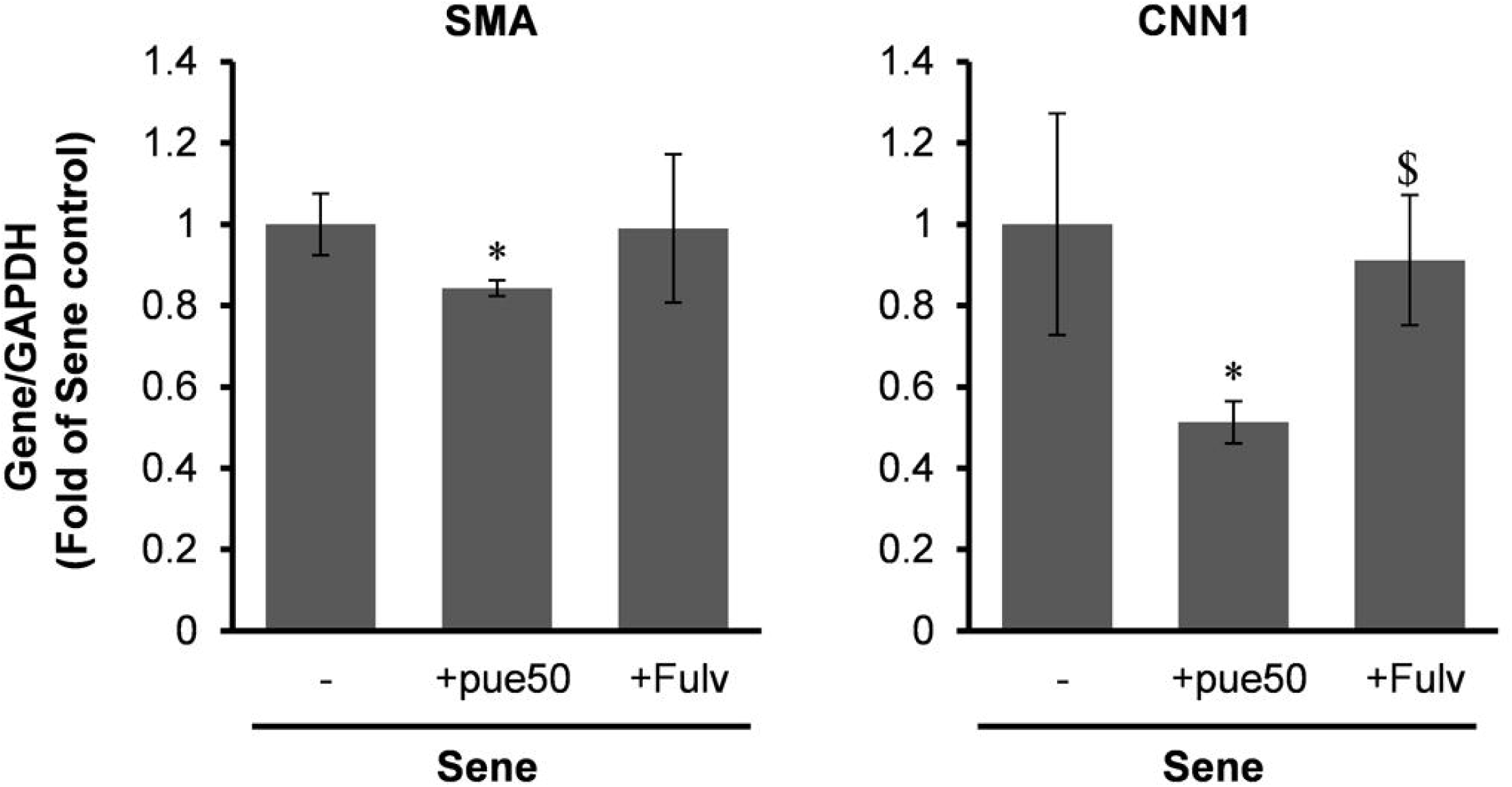
Estrogen receptor antagonist blocked puerarin-mediated reduction of SMA and CNN1. The expressions of SMA and CNN1 mRNAs in senescent NHDFs treated with 50 μM puerarin (+pue50) and estrogen receptor antagonist, 1 μM fulvostrant (+Fulv) for 72 h. n=4 *; p<0.05, vs control, and $; p<0.05 vs +pue50.

## Discussion

In the present study, we evaluated the effect of puerarin in the senescent NHDFs and discovered that puerarin enhances the viability of NHDFs accompanied by the increased proliferative activity and the decreased cell injury. Furthermore, we found that puerarin attenuates the frequency of SA-beta-galactosidase positive senescent cells. Puerarin also attenuates aging-related fibroblast subpopulation changes; i.e. a decrease in myofibroblasts and reticular fibroblasts by puerarin. Importantly, high passages cause to differentiate from papillary fibroblasts into reticular fibroblasts, suggesting that puerarin may prevent age-related fibroblast transdifferentiation.

While puerarin possesses a wide variety of pharmacological properties, various molecular mechanisms have also been proposed [29]. Among them, puerarin is recognized as a phytoestrogen. Puerarin or the extract of *Pueraria Lobata* has a beneficial effects on menopausal symptoms, such as bone loss [25]. Whereas puerarin has a weak estrogenic activity, puerarin also exerts an antagonistic effect against 17-β-estradiol and estrogen-response element-reporter transcriptional activity [29]. In the present study, we found that ER inhibitor, Fulvestrant, blocked the effect of puerarin on the SMA and CNN1 expressions, indicating that the effects of puerarin on senescent NHDFs are via ER pathway. Since estrogens are key hormones for skin aging homeostasis, and act directly on dermal fibroblasts [43,44], puerarin might contribute to this mechanism.

Additionally, puerarin possesses antioxidant properties [29]. In the present study, we found that ROS production was not decreased, rather increased in puerarin-treated NHDFs. In the case of proliferation-induced senescent cells, excessive ROS production may not be associated with cellular defects. In contrast, the appropriate generation of ROS has beneficial anti-aging roles by modulating some signaling pathways [2]. Therefore, puerarin-induced ROS generation may be involved in its anti-aging effect in NHDFs.

In the present study, we observed a reduction in the frequency of SA-β-gal positive cells by puerarin, suggesting the senolytic property of puerarin in NHDFs. However, we also observed that puerarin did not cause to induce cell toxicity in high-passaged NHDFs, as confirmed by LDH assay and the ethidium homodimer-I assay (live-dead assay, data not shown). Taken together, puerarin is not as a senolytics for senescent NHDFs.

Our present study indicates that the *Pueraria lobata* diet may confer anti-aging properties in the skin. Meanwhile, the bioavailability of orally ingested puerarin has been evaluated in rodents, and it is approximately 4-7% [34]. When 10 mg/ kg body weight of puerarin was administered to the rodents, Cmax of blood puerarin was 228 μg/L (approximately 0.5 μmol/L). In the present study, we confirmed the short-term effect of puerarin at 25 to 50 μmol/L in the culture medium, and it is much higher than the blood concentration expected to be after the *Pueraria lobata* diet. Long-term and low-dose *Pueraria lobata* diet study will be necessary to confirm the clinical effect in the future.

## Abbreviations

NHDFs: normal human dermal fibroblast
SMA: smooth muscle actin
CNN1: calponin 1
ER: estrogen receptor

## Acknowledgements

Funding: This work is supported by the grant for Scientific Research from Fukui Prefectural University.

